## Supplementary figures and images for "Plant-Necrotroph Co-transcriptome Networks Illuminate a Metabolic Battlefield"

### Supplemental Figure 1-8

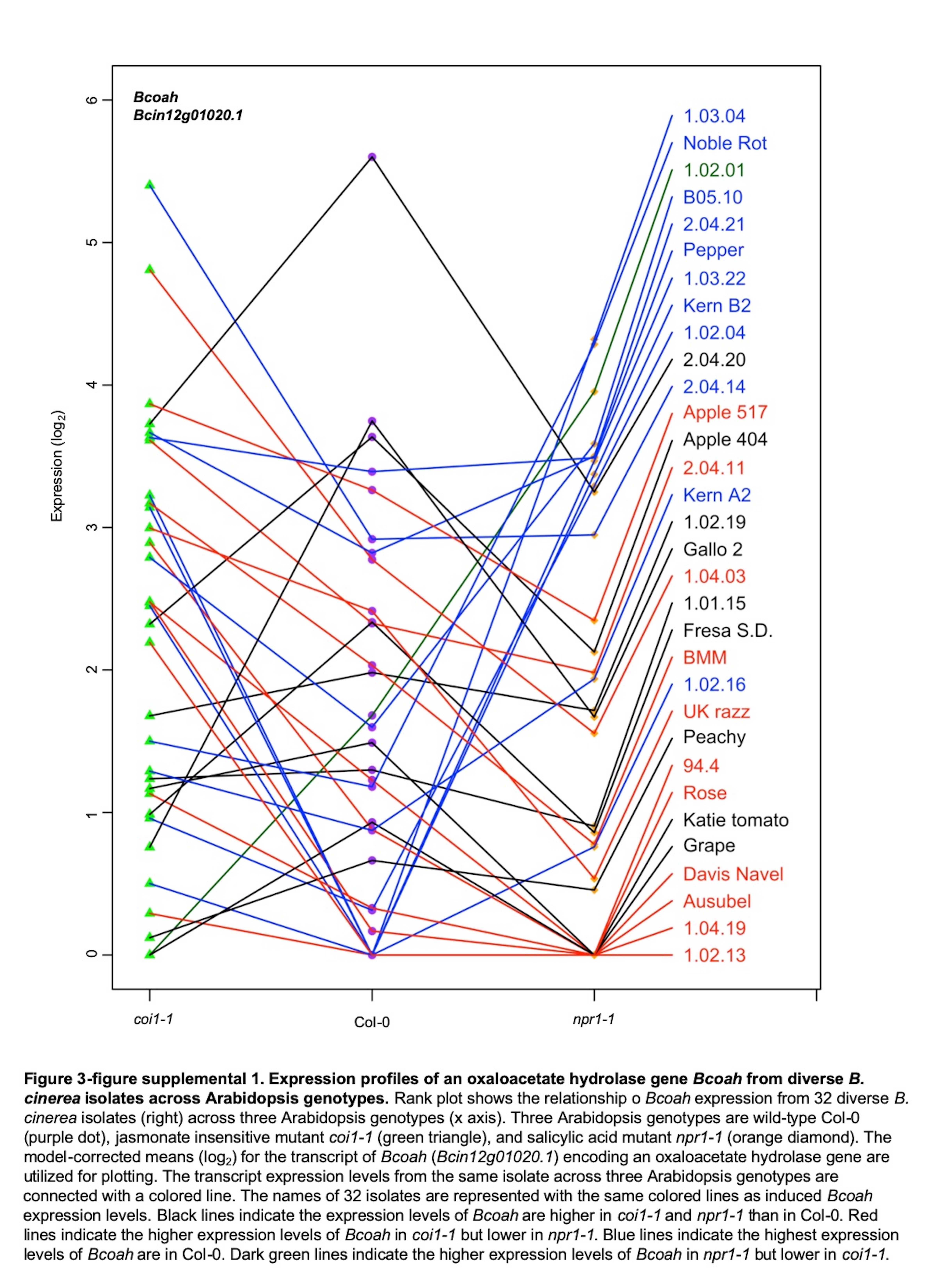

### Supplemental Figure 1-8

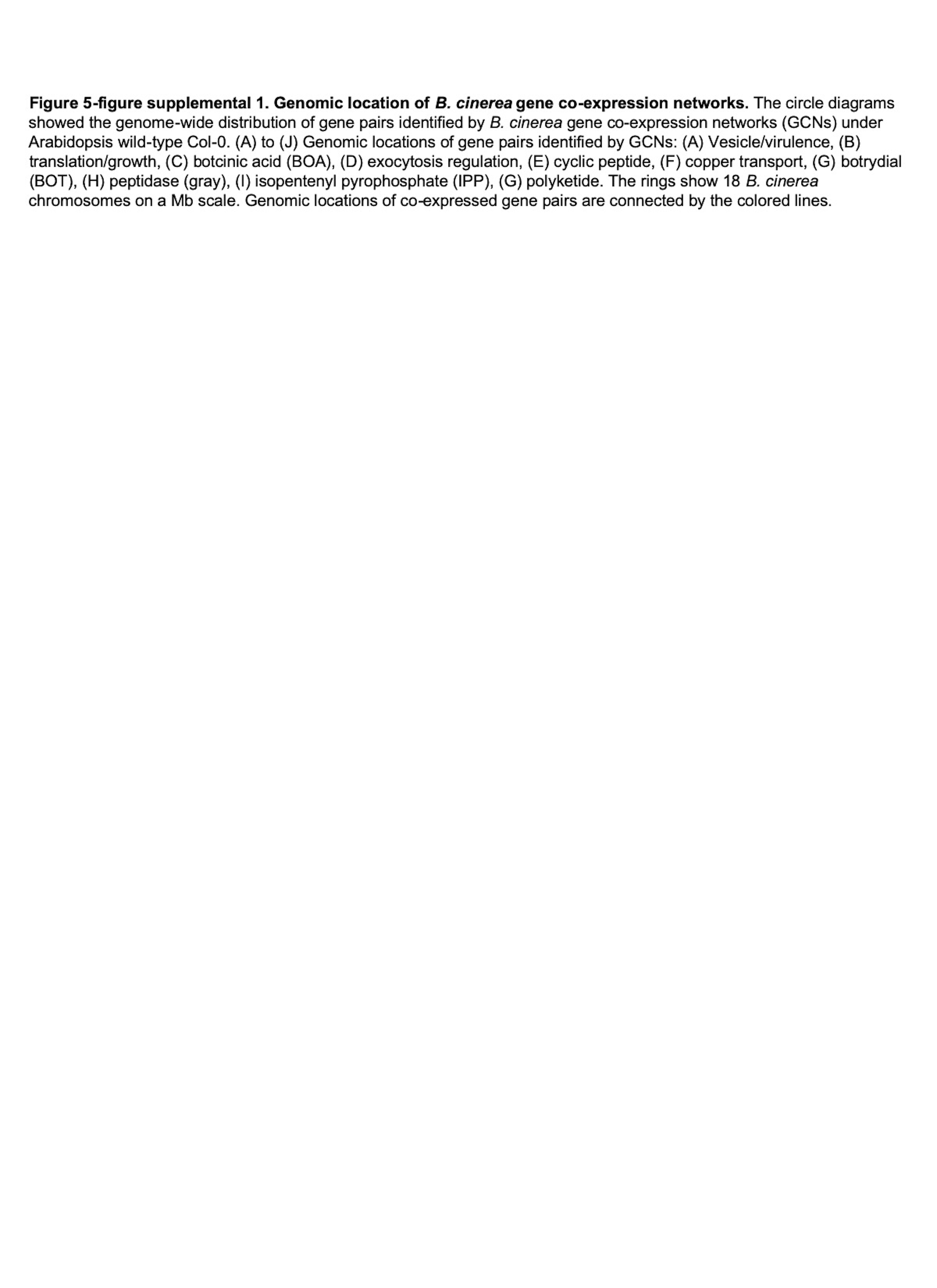

### Supplemental Figure 1-8

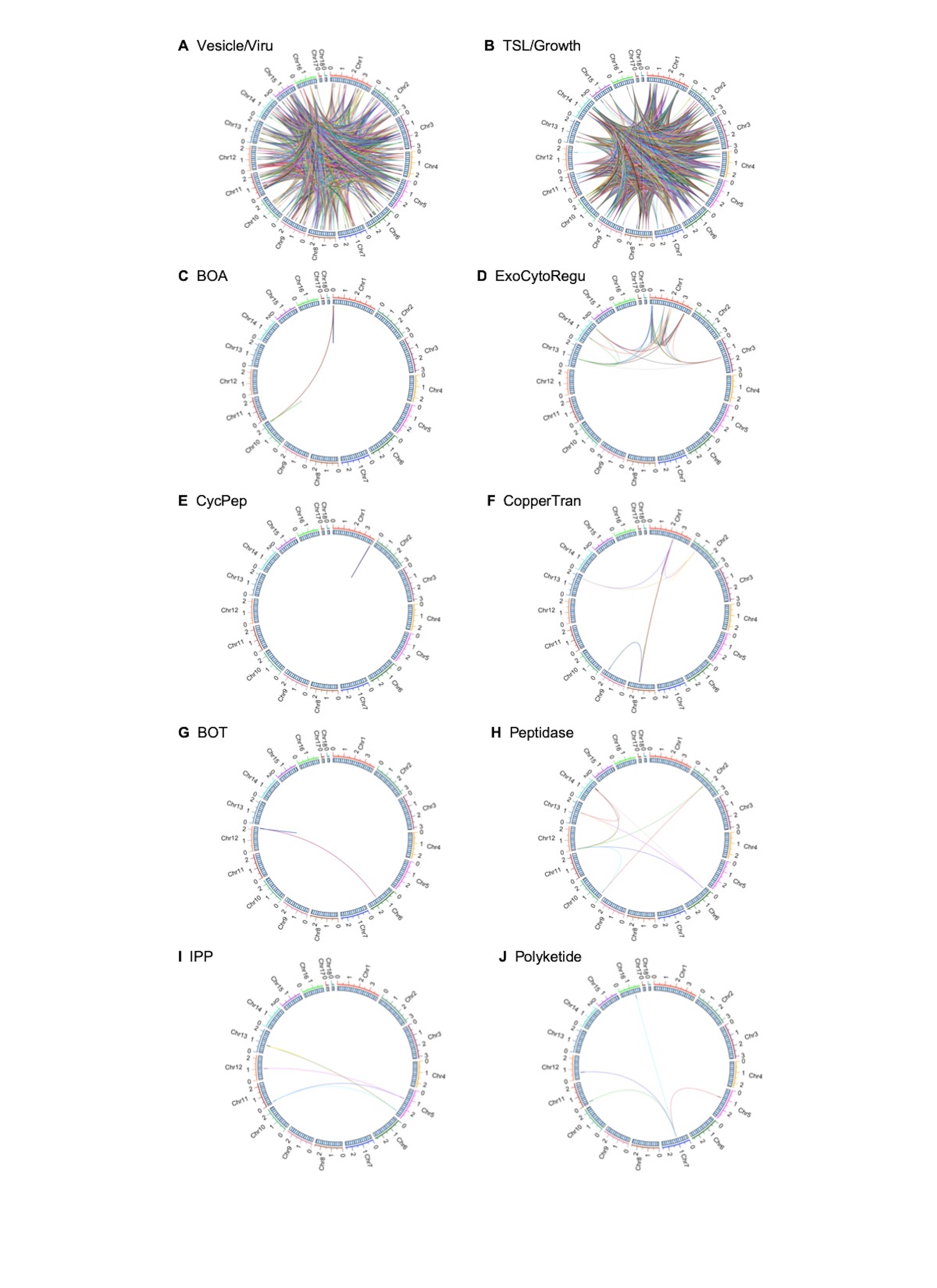

### Supplemental Figure 1-8

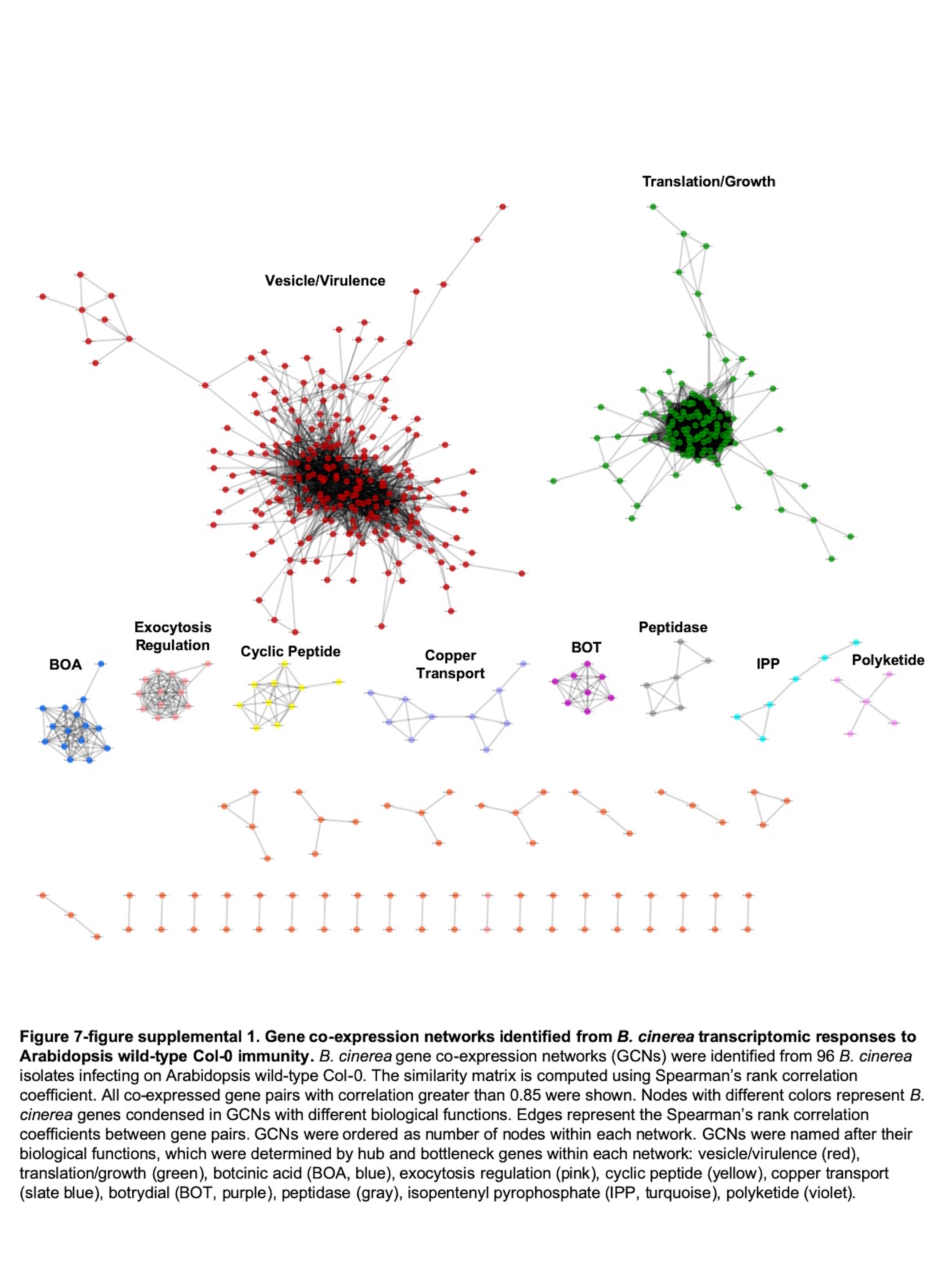

### Supplemental Figure 1-8

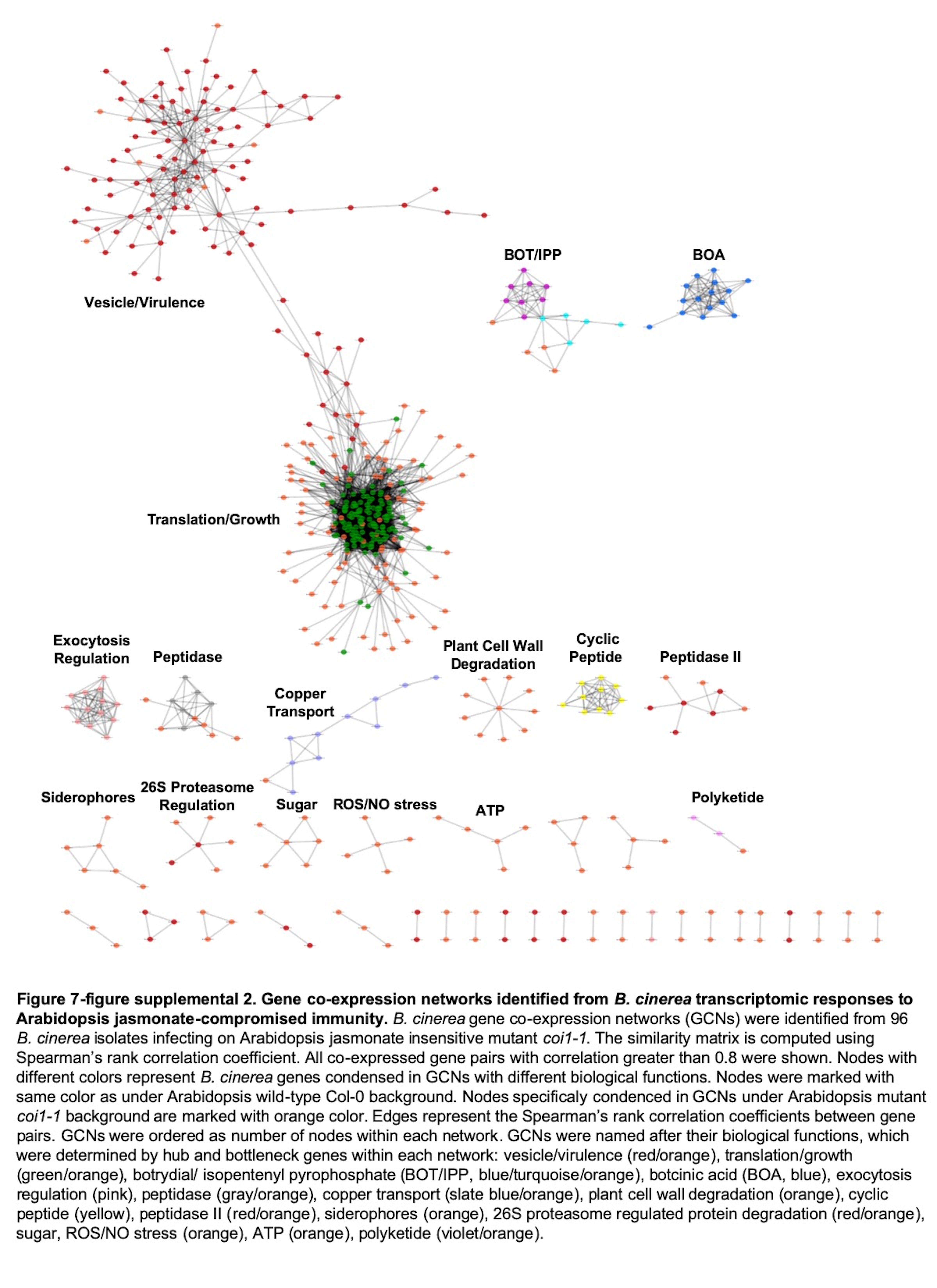

### Supplemental Figure 1-8

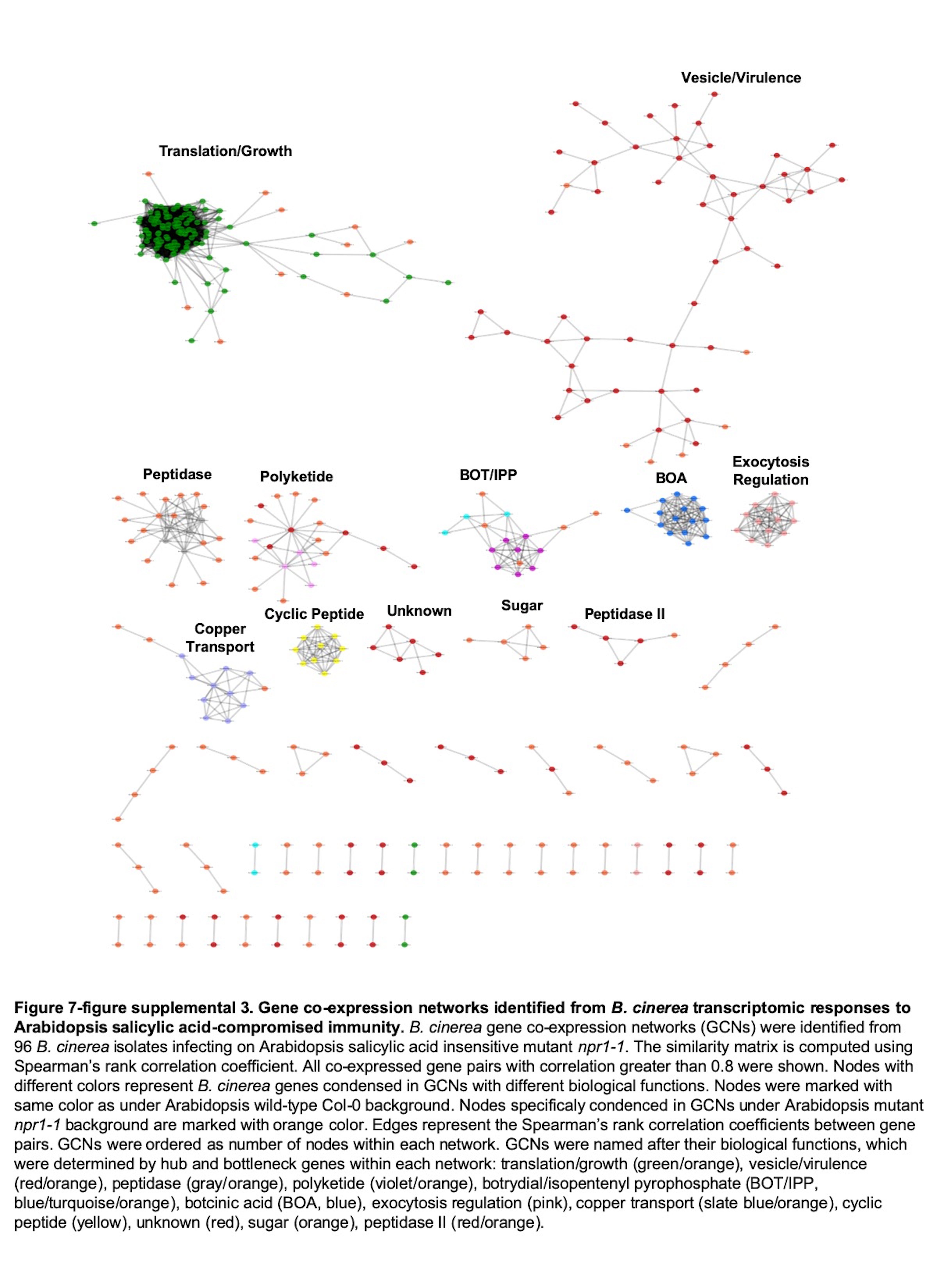

### Supplemental Figure 1-8

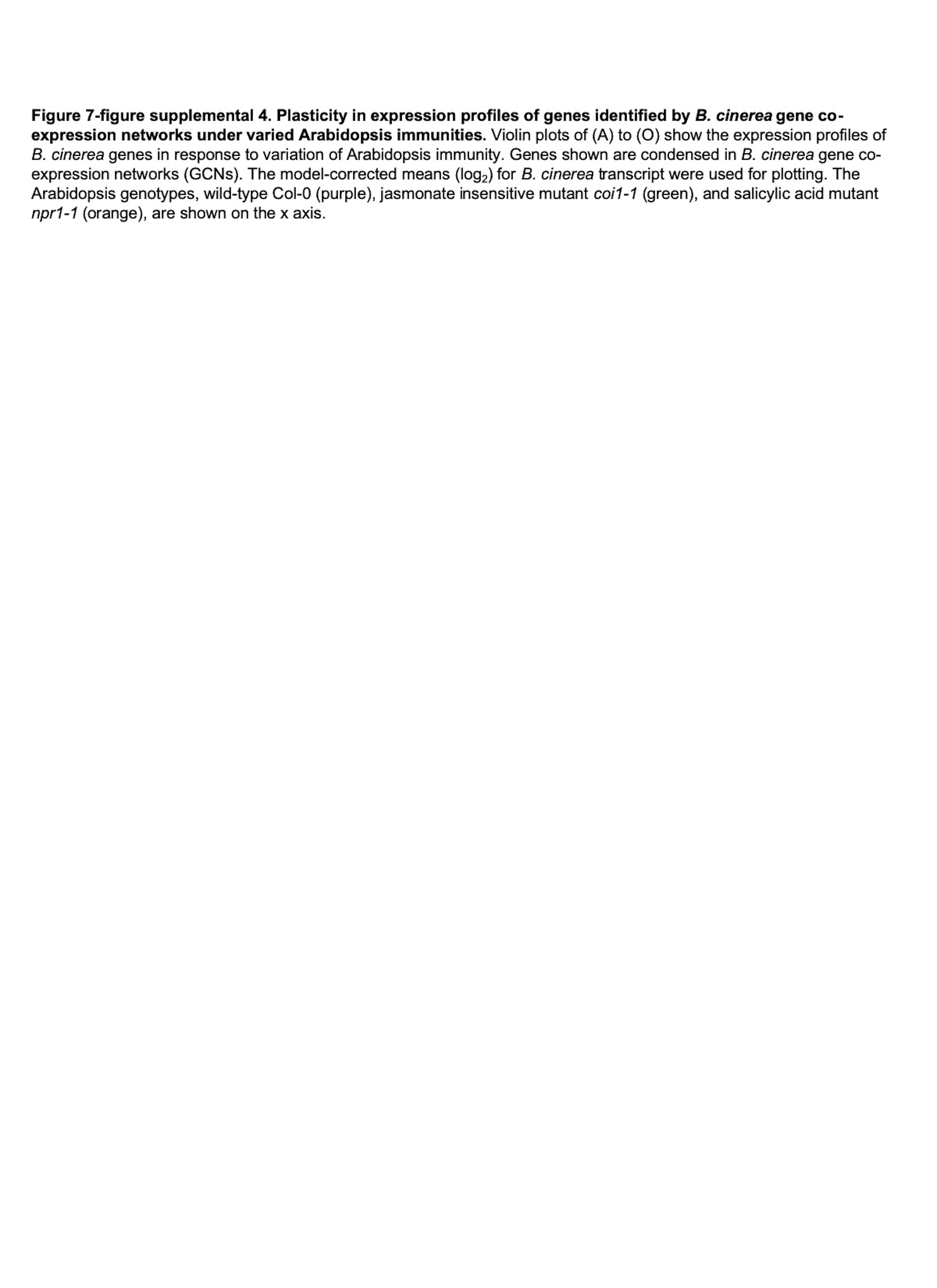

### Supplemental Figure 1-8

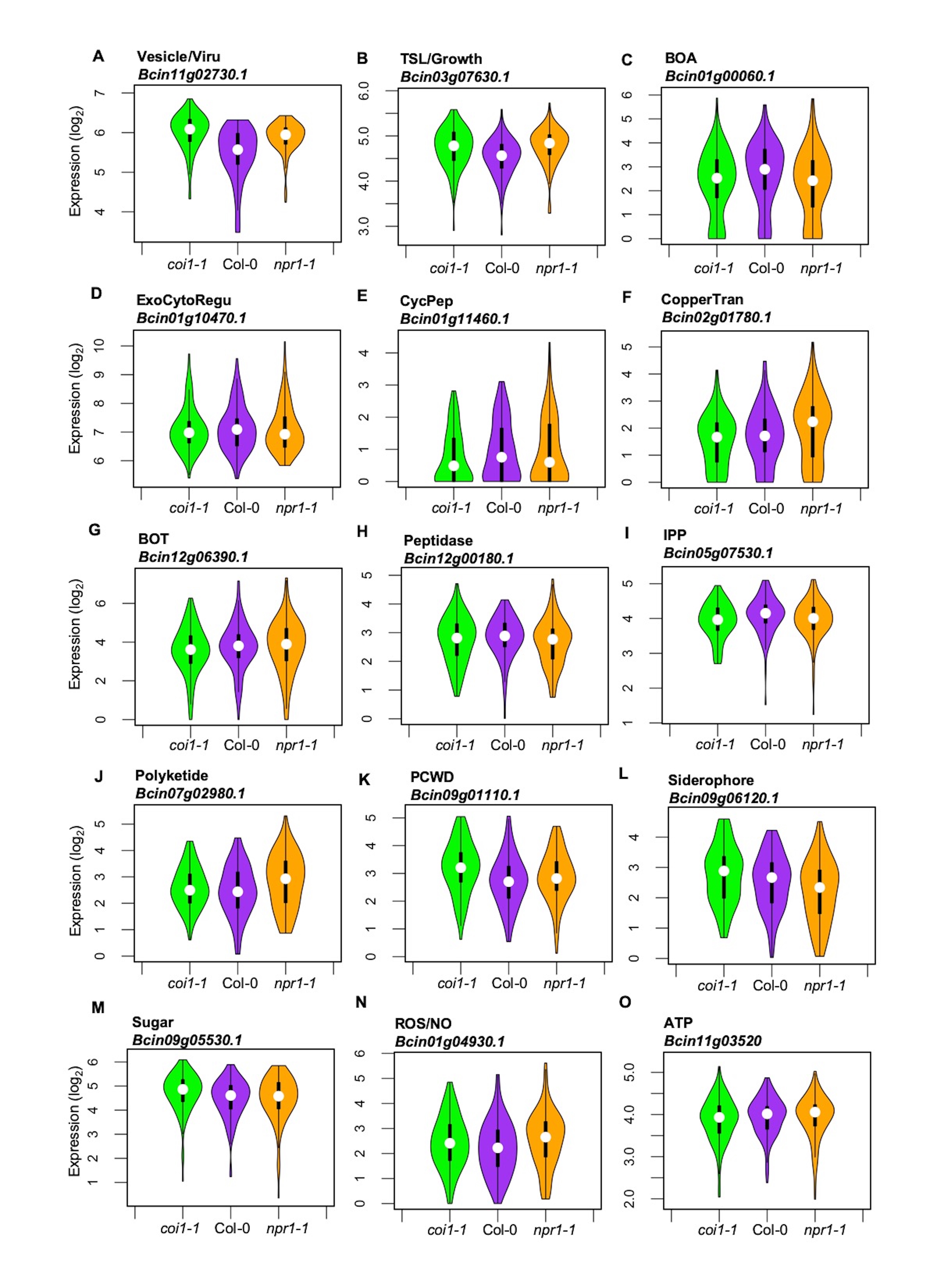

### Supplemental Figure 1-8

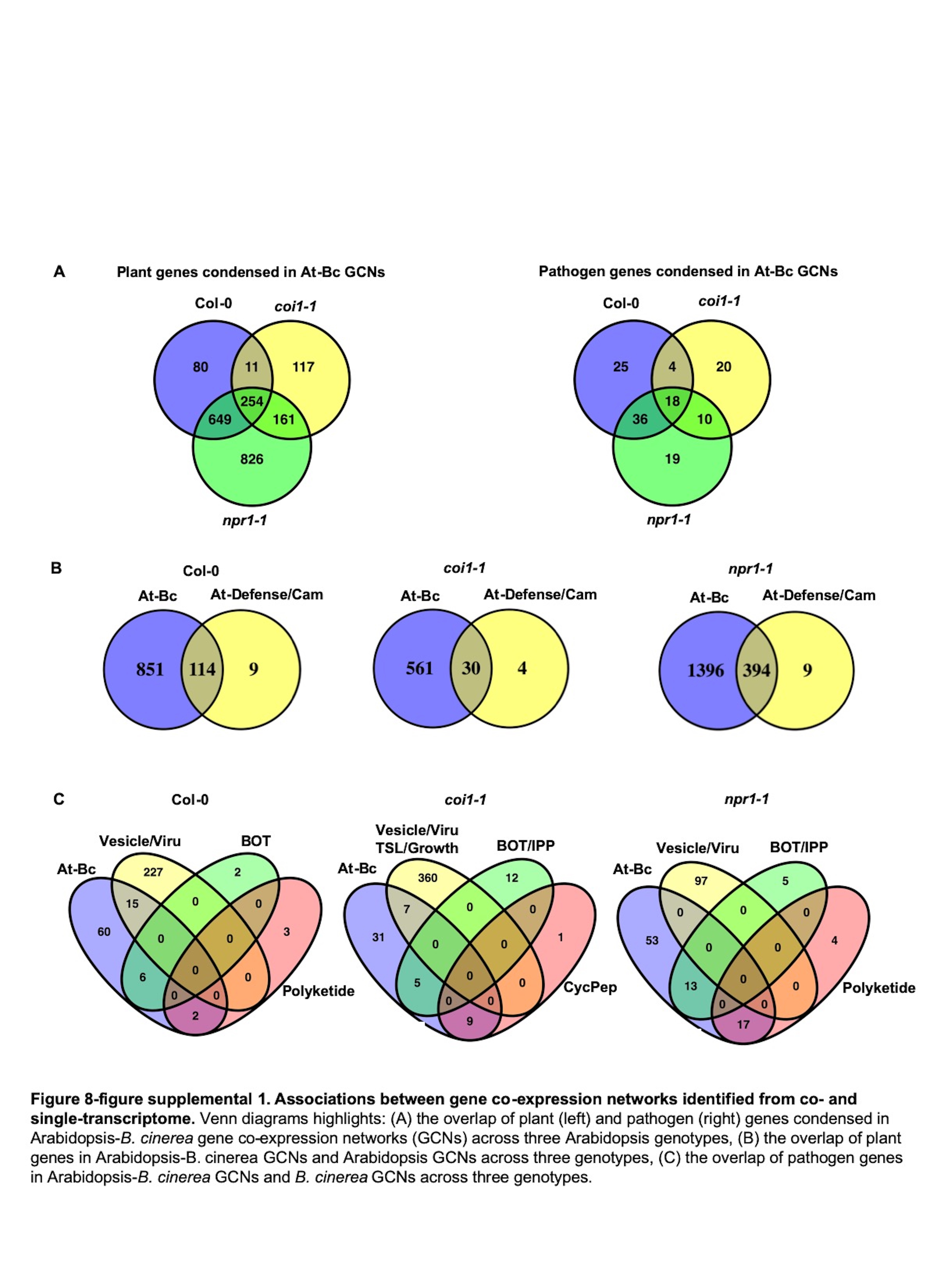

### Supplemental Figure 1-8

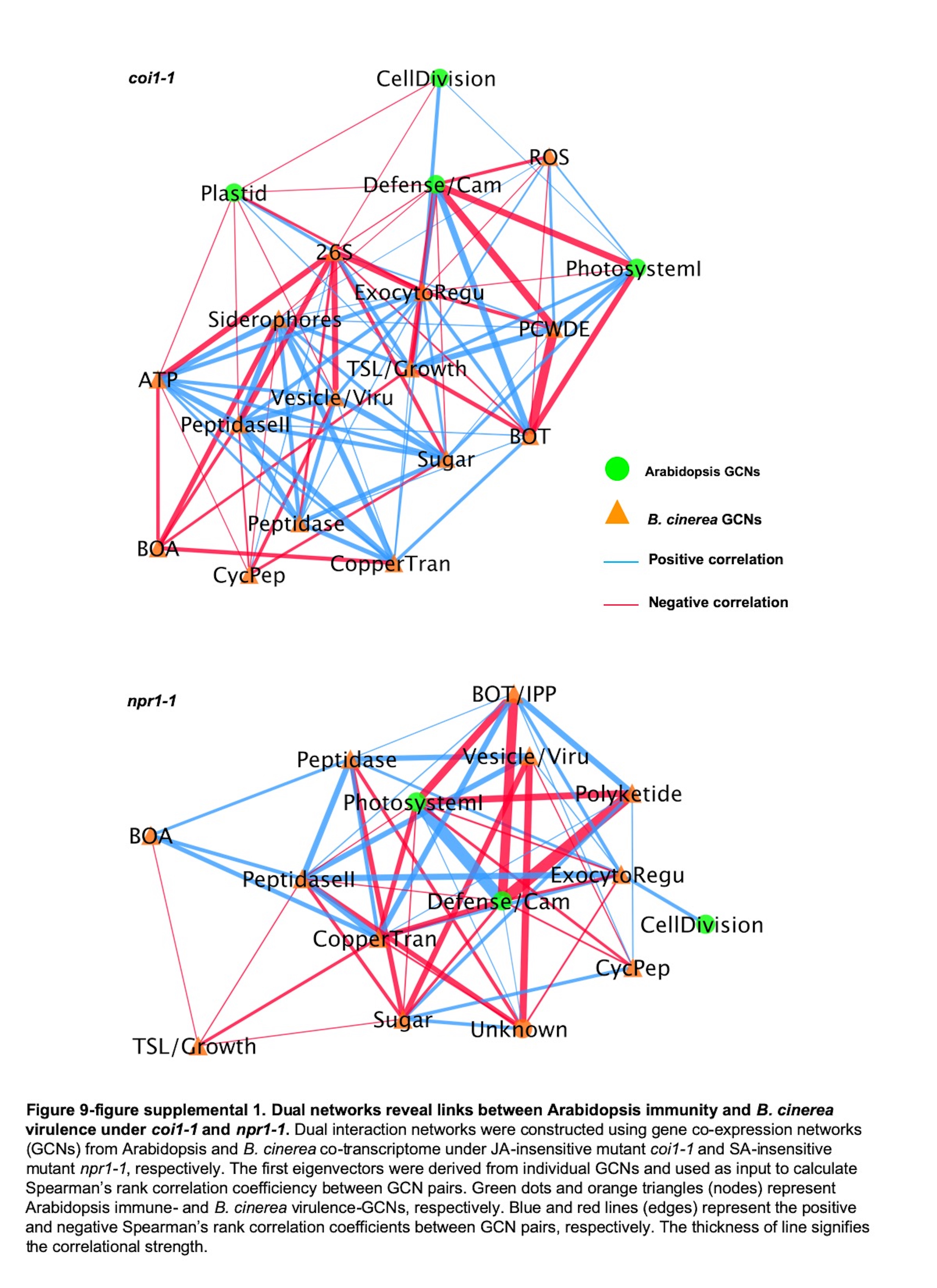

### Supplemental Figure 1-8

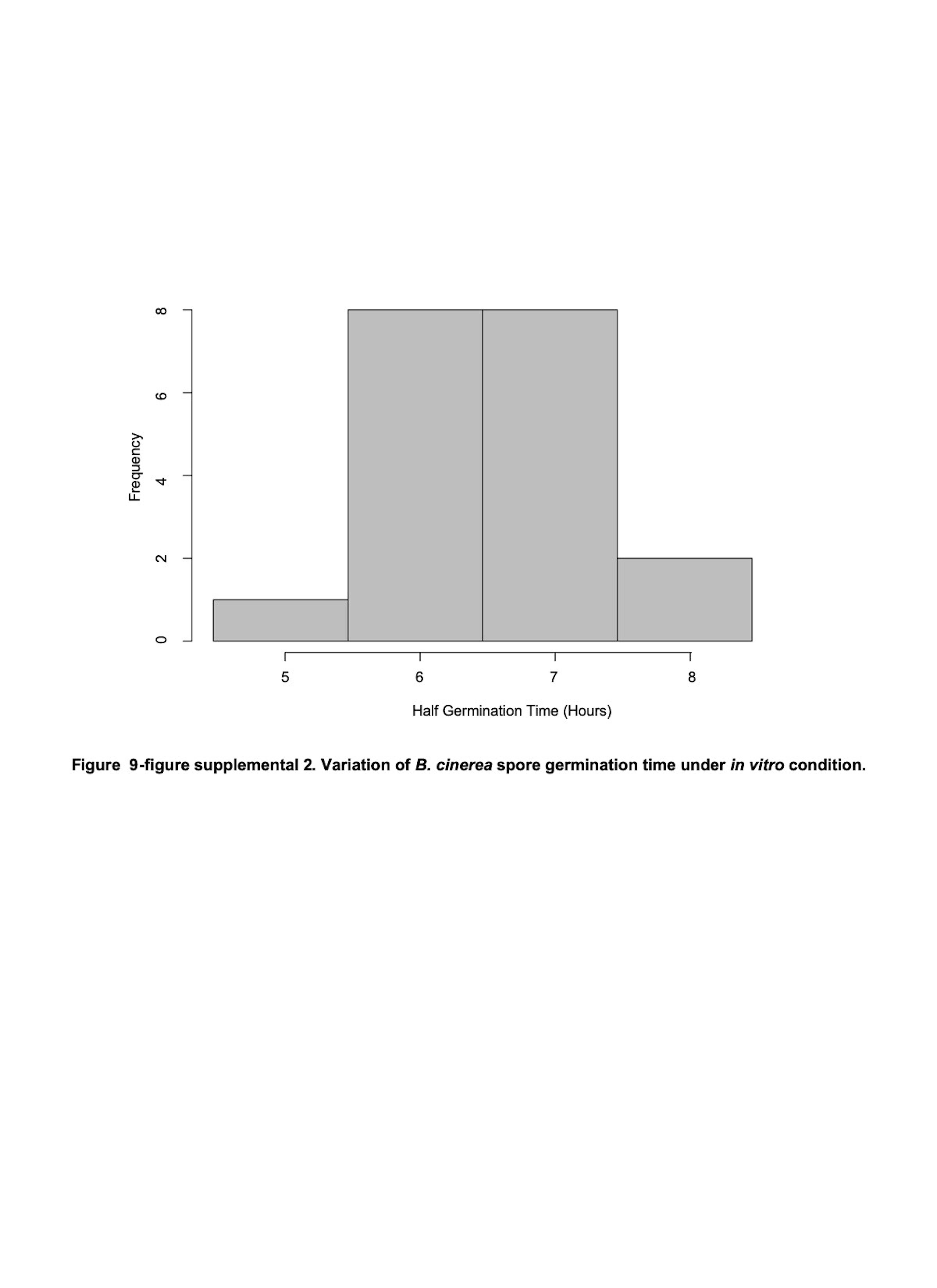
